# A touchscreen device for behavioral testing in pigs

**DOI:** 10.1101/2022.08.18.504438

**Authors:** Will Ao, Megan Grace, Candace L. Floyd, Cole Vonder Haar

## Abstract

Pigs are becoming more common research models due to their utility in studying neurological conditions such as traumatic brain injury, Alzheimer’s disease, and Huntington’s Disease. However, behavioral tasks often require a large apparatus and are not automated, which may disinterest researchers in using important functional measures. To address this, we developed a touchscreen that pigs could be trained on for behavioral testing. A rack-mounted touchscreen monitor was placed in an enclosed, weighted audio rack. A pellet dispenser was operated by a radio frequency transceiver to deliver fruit-flavored sugar pellets from across the testing room. Programs were custom written in Python and executed by a Raspberry Pi. A behavioral shaping program was designed to train pigs to interact with the screen and setup responses for future tasks. Pigs rapidly learned to interact with the screen. To demonstrate efficacy in more complex behavior, two pigs were trained on a delay discounting tasks and two pigs on a color discrimination task. The device held up to repeated testing of large pigs and could be adjusted to the height of minipigs. The device can be easily recreated and constructed at a relatively low cost. Research topics ranging from brain injury to pharmacology to vision could benefit from behavioral tasks designed to specifically interrogate relevant function. More work will be needed to develop tests which are of specific relevance to these disciplines.

## 1. INTRODUCTION

Historically, the rodent has been the model of choice for neuroscience research. There are several reasons for this, including the economical nature, both in cost and space compared to larger species. As such, an abundance of methods and tools have been developed for use in rodents. Behavioral assessment is particularly well-developed with a number of sensorimotor tests and more complex tasks which measure human-relevant functions such as decisionmaking, working memory, and self-control. However, there are also concerns with rodent models. The physiology is dissimilar to humans, particularly the brain structure and size. To effectively develop a translational pipeline for neurotherapeutics and to understand neuropathology, larger species with greater brain similarities to humans are needed.

One such existing model is the pig, already well-established for the study of circulatory, nervous, and respiratory function due to their physiological similarities to the human [1]. Pigs are especially attractive in neuroscience because of a high degree of similarity to human brains in the sulci and gyri, with gyrification far exceeding the rodent model and even exceeding a common non-human primate model, the rhesus macaque [2]. Given the greater anatomical homology, CNS diseases, injuries, and challenges in pigs are much more likely to cause pathology of greater similarity to humans relative to rodent models. For example, the pig cortex is more compressible than rodents [3]. Pig brains are also considerably larger than their rodent counterparts, weighing in at 95.3 g relative to less than 2.5 g in the case of the rat. This size again exceeds the rhesus macaque (90 g) [2].

Recently, several fields have increased the use of pig models. Pigs have become prominent in the field of traumatic brain injury (TBI), where they serve multiple purposes such as analyzing pathophysiology, understanding surgical management of injury, and even studies on recovery of function [4]. In addition, the recent development of transgenic Göttingen minipigs have created a strong model for studying the pathology of Alzheimer’s disease (AD; amyloid precursor protein and *presenilin-1* mutations) and Huntington’s disease (HD; *huntingtin* mutation) [5, 6], while a transgenic of the Minnesota minipig has been created for cancer research (floxed line for cre-dependent tumor expression) [7]. Notably, each of these conditions strongly impact behavioral function, however porcine cognitive assays are relatively limited. Simple discriminations, reversals, and working memory tasks have been used in the case of TBI [8, 9]. Motivation or ability to uncover hidden treats has been used in the HD transgenic model [10] and a recognition memory task (novel object test) in the AD transgenic model [11]. However, even these tasks do not always distinguish the condition from control and small effect sizes of injury or disease may limit detection of treatment effects. In surveying this literature, we have identified three primary barriers to expanding cognitive testing in pigs: 1) specialized equipment requirements, 2) behavioral expertise of the experimenters, and 3) study time constraints. While item number 3 will be inherently study dependent, the first two challenges can be addressed to some degree with technology.

Historically, both rodents and primates had purpose-designed behavioral equipment. Perhaps most notable is the operant chamber, which is a modular, computer-controlled chamber in which many different cognitive tests can be assessed and has a small space footprint. In contrast, current functional assays for pig behavior often require large, room-sized apparatus and manual setup and scoring. Even traditional T-mazes or multi-arm mazes become a challenge due to the space constraints and are inherently low throughput. Manual tasks also introduce the possibility of unconscious experimenter bias. While uncommon, several researchers have adapted automated devices (including a primate operant chamber) for testing cognitive function in pigs. These devices have been used to assess complex cognitive behaviors such as behavioral inhibition [12], working memory [13], and behavioral flexibility [14]. Others have set up similarly sophisticated behavioral assays, including gambling-like behavior [15], choice impulsivity [16], and working memory [17] but required full manual administration and/or room-sized mazes. The heterogeneity in testing apparatus and manual nature of many are likely a leading reason for sparse adoption of pig cognitive outcomes. As such, the development of a small footprint, low-cost platform with the capability to perform high throughput operant measures is needed.

To develop such a device, we can make use of touchscreen technology, open-source software, and readily available equipment. This may provide the benefit of standardizing methods across species, including humans. A recent argument has been made that this will help close the gaps in fields that have struggled with translation, such as pharmacology [18], although task similarity is likely a more important component than test medium (e.g., touchscreen). In the current paper, we describe the development of a device to make behavioral research for the porcine model more accessible to a wide variety of researchers. We utilized relatively low-cost materials and open-source software to create a robust tool for behavioral analyses. This can be constructed in the average laboratory environment by ordering the commercial pieces and assembling or can be made even lower cost with some modifications noted in the methods.

## 2. MATERIALS AND METHODS

### 2.1 Touchscreen Design Overview

The overall device design and finished product is shown in Figure 1. Two versions were constructed; the first iteration was susceptible to damage from the pigs and is only briefly described in the results for transparency (but can be viewed in video S1). Subsequent methods will refer to the more durable, second iteration. A touchscreen monitor was attached to a Raspberry Pi which ran custom behavioral programs. In this final iteration, a radio frequency (RF) transceiver was attached to the input/output pins of the Raspberry Pi. This allowed pellet delivery to be located across the room to reduce rooting behavior toward the screen. The RF transceiver communicated to a second Raspberry Pi with a receiver. The second Raspberry Pi input/output pins were hooked up to a printed circuit board (PCB). The PCB contained 8 inputs and 8 outputs capable of handling standard 28V operant equipment. In the iteration described here, only a pellet dispenser and Sonalert tone generator were attached as outputs.

**Figure 1.**
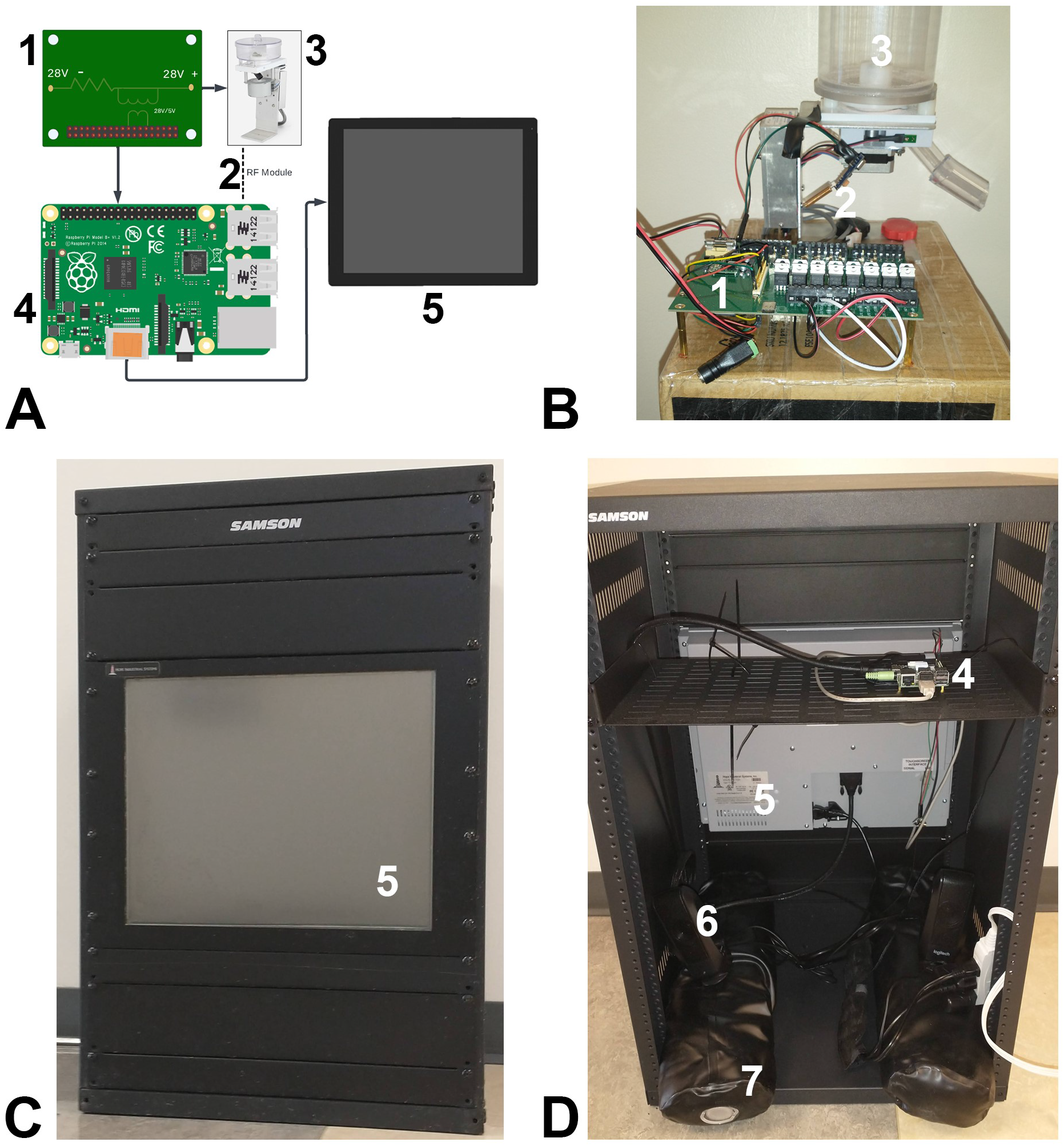
Schematic layout and actual device. A) The conceptual schematic organization of the touchscreen device with pieces numbered in black, corresponding to white numbers on the actual device (panels B-D). A Raspberry Pi controls the touchscreen and records responses. Output is then taken from the Raspberry Pi I/O pins and put through an external printed circuit board to step the voltage up to the 28 V needed for peripheral components. Physical inputs were available but not used in the current studies. B) An image of the remote-controlled dispenser, attached to the PCB. A raspberry Pi is attached underneath the PCB and not visible. Not in this picture is the sonalert tone generator which was also hooked up to the PCB. C) Front of testing device showing screen. D) Rear of testing device showing back of screen, Pi, and other components. Numbered items indicate: 1) PCB, 2) RF transceiver, 3) pellet dispenser, 4) Raspberry Pi, 5) touchscreen, 6) audio speakers, 7) weighted sandbags.

### 2.2 Physical Components and Construction

All components are detailed in Table 1 along with the cost to acquire at time of purchase. The below description is for the final iteration of the device.

**Table 1.** List of physical components and pricing as of purchase. Sums in each category are given in bold. *The board and components were purchased as a set of 5 to meet minimum order thresholds; price given is per board. **A smaller rack would have been suitable even for the large Yorkshire pigs. Future builds will use the SRK-12 (12U height). ***The rack shelf was not explicitly necessary, but gave easy access to the Raspberry Pi.

| Item | Parts | Price | Vendor |
| --- | --- | --- | --- |
| **Computer** | | **\$148** | |
| | Raspberry Pi 4B (2x) | \$110 | SparkFun |
| | Power supply (2x) | \$16 | SparkFun |
| | Misc. jumpers etc. | \$10 | SparkFun |
| | RF Transceivers | \$12 | Amazon |
| **PCB** | | **\$39** | |
| | Board* | \$19 | PCBWay |
| | Misc. components* | \$20 | Digikey |
| **Peripherals** | | **\$799** | |
| | Pellet Dispenser | \$729 | Med-Associates |
| | Sonalert | \$70 | Med-Associates |
| | Desktop speakers | \$0 | in lab |
| **Monitor** | | **\$855** | |
| | 19" Rack-mounted Resistive | \$835 | Hope Industrial |
| | Screen protector | \$20 | Hope Industrial |
| **Frame** | | **\$238** | |
| | Samson SRK-16** | \$165 | B&H Audio-Video |
| | 1U blank panel (4x) | \$18 | B&H Audio-Video |
| | 2U blank panels (2x) | \$12 | B&H Audio-Video |
| | rack shelf*** | \$15 | B&H Audio-Video |
| | Sandbags (2x) | \$28 | Amazon |
| **Total** | | **\$2,079** | |

#### Frame

Standard 19” wide racking for servers or audio equipment was used to house the screen. It was enclosed on two sides and 31” high (16U rack height), 18” deep. Rack “blanks” were used to fill the front not occupied by the screen and the rear was left open but put against a wall. A rack -mounted shelf was put in the rear to hold the Raspberry Pi. The modular nature of the frame allowed the screen to be moved up or down to deal with smaller or larger pigs. A pair of sandbags (9 kg each) were purchased and filled to weigh down the device so that pigs could not move it.

#### Screen

A rack-mounted resistive touchscreen (Hope Industrial Systems, HIS-RL19-CTDH) designed for industrial use was mounted and adjusted for the height of the pigs. A plastic screen protector was placed over the screen to reduce scratches and for easier cleaning.

#### Circuit Board Interface

A PCB was custom-designed to take 28V input/output to 3.3V for interfacing to the Raspberry Pi. For outputs an optical switch was used to isolate the 3.3V from 28V. For inputs, resistors were used to step down the voltage. A pass-through so that the board could power the Pi was used.

#### RF Transmitter/Receiver

A simple 433 MHz radio frequency transceiver/receiver combination were purchased, and an antenna soldered to each. Software was modified to turn received signals into outputs (pellet, tone) on the receiving Raspberry Pi and to transmit them on the sending Raspberry Pi. The pellet dispenser was attached to a box which sat on top of a sink, raising it above the reach of the pigs.

#### Raspberry Pi

Two Raspberry Pi model 4B were used, along with power supplies and jumper cables to connect.

#### Peripheral Components

A pair of desktop computer speakers were placed in the bottom of the frame to provide auditory feedback. A pellet dispenser (Med Associates, ENV-203-1000) was connected to the receiving raspberry pi via the PCB, located across the room. A Sonalert tone generator (Med-Associates ENV-223AM; 2900 Hz, 100 dB) was also attached alongside the pellet dispenser.

#### Alternative Materials

The largest costs in construction are the monitor, pellet dispenser, and frame. Cheaper framing materials may be available from other vendors, although we recommend a reputable brand that will stand up to pigs interacting with it. Other rack-mounted screen options are also available, although price points vary widely. We recommend choosing one that has a reasonable impact rating and does not protrude from the racking as pigs may chew at the corners. For the dispenser, it is possible to 3D print and purchase small motors to operate it as described in papers for rats [19, 20]. Alternate food delivery systems would likely also be suitable.

### 2.3 Software Components

#### Peripheral Device Control

A program to send and receive RF signals in Linux was modified from RFChat [21]. A program was written and ran on boot of the receiving Raspberry Pi. On receipt of a given cue (e.g., “1”), the pellet dispenser would cycle. On receipt of another cue (e.g., “2”), the tone would turn on for 1 s. Thus, as long as the board attached to the receiving Raspberry Pi was powered, it would control the peripheral devices. Behavioral programs on the sending Raspberry Pi used these commands on relevant events (i.e., reinforcement).

#### Graphical User Interface (GUI)

The Python Tkinter package was used to develop GUIs. For human user input, a pop-over box with options to change variables (e.g., subject number, training stage, etc.) in the underlying behavioral program populated at the start. A program for a touchscreen numpad was written to allow numbers to be input without a keyboard (a wireless keyboard may also be used). For pig responses, buttons housed within a full screen window were used. Buttons were designated to respond on initial touch (default is release of click or removal of touch).

#### Data Recording

Every response was recorded as a new line in a comma-separated value file with a separate file for each subject. Each behavioral program recorded information relating to the individual trial as well. Summary data were reported on the screen at the end for daily monitoring of progress.

#### Behavioral Testing Programs

Custom programs were written in Python according to the descriptions below. A common shaping program was used to train initial response to the screen and to smaller boxes within the screen.

#### Data Transfer To/From Raspberry Pi

An FTP server (vsftpd package) was setup on the raspberry pi with a folder to which a remote computer could read and write files. An FTP transfer utility (FileZilla) was used to move behavioral programs onto the Pi and pull data files for analysis.

### 2.4 Evaluation of Device Durability

The core goal of the current studies was to determine if a touchscreen device would withstand repeated testing with pigs. In our first iteration of the device, the screen was broken after 28 sessions with two pigs. In the second iteration, which is what is described above, the screen withstood testing throughout all 35 sessions and remains intact.

### 2.5 Subjects

Male (castrated) and female Yorkshire pigs were used in the described experiments. Two male pigs were 9 weeks of age at start of training, weighed 24-26 kg and were tested in experiment 1. Two female pigs were 12 weeks of age at start of training, weighed 45-52 kg and were tested in experiment 2. Pigs were obtained from the Ohio State University farms and were acclimated to the vivarium for one week prior to testing. Pigs were housed in individual pens adjacent to one another. Males and females were housed separately. All procedures were approved by the Ohio State University Institutional Animal Care and Use Committee.

### 2.6 Behavioral Training

#### Reinforcers

Mini-marshmallows were used as reinforcers during pre-training and 1 g fruit-flavored sucrose pellets (Bio-Serv F05478, F05711) were delivered from the pellet dispenser while pigs interacted with the touchscreen.

#### Pre-Training of Pigs

The goal of this step was to familiarize pigs with the experimenter, get them used to leash walking, and traveling to the testing room using basic behavior shaping techniques with a clicker. In their home room, pigs were trained to associate a clicker with mini marshmallow delivery and approach the experimenter to receive the reinforcer. The experimenter then familiarized them with the leash by draping it across them, then wrapping it around them, while providing reinforcers. Once comfortable, a large dog harness was placed over the shoulders and clipped behind the legs. Pigs were then trained to walk on the lead in the home room while receiving reinforcers. Once comfortable, the outer hallway was blocked off (either physical blockade, or second researcher with a board) and pigs were taken back and forth down the hallway. Once pigs were responding well to the leash, they were led to the behavior testing room. This process could be accelerated with multiple sessions per day if needed.

#### Response Shaping

To shape responses to the touchscreen, a multistage procedure was followed. Pigs moved up in stages automatically within the program or were started at later stages if they had completed the prerequisite the session before. Audio speakers inside the device provided auditory feedback when the button was pressed (2900 Hz tone) and when a trial began (7500 Hz tone).

Stage 0 was a Pavlovian autoshaping procedure in which the entire screen illuminated (yellow color) and then a pellet was delivered 10 s following illumination. However, there was also a fixed ratio (FR)-1 schedule in effect such that at any given time, a press to anywhere on the screen would be reinforced. If a pig did not contact the screen, ketchup was wiped on the screen to motivate approach. After 20 presses, pigs moved to stage 1. Stage 1 made the response conditional - presses were only reinforced when the screen was illuminated (FR-1 schedule). After 15 presses, pigs moved to stage 2. Stage 2 reduced the size of the response box from the entire screen to a large, illuminated (yellow) rectangle occupying 1/3 of the screen. The rectangle was positioned randomly at one of three heights to shape responses to track the change in position. Responses to the rectangle were reinforced (FR-1 schedule). After 15 presses, pigs moved to stage 3. Stage 3 reduced the size of the response box further to a square (yellow, 40% of screen width) which was positioned randomly in one of five positions (just offset from each corner, and center). After 40 presses, pigs were considered ready for testing. A few optional manipulations may be considered at this point. If a higher response requirement will be needed for subsequent tasks, stage 3 may be increased to FR-3 or FR-5. If smaller response boxes will be needed, a stage 4 where the box size shrinks gradually over successive correct trials may help shape precision. Close attention should be paid throughout to the behavioral topology, or the way in which responses are made and how that may affect subsequent tasks. See results for qualitative descriptions of pig behavioral topology in interacting with the device.

### 2.7 Behavioral Assessment of Pigs

After shaping nose pokes to illuminated buttons, pigs were tested in one of two experiments. Experiment 1 used two male pigs, while Experiment 2 used two female pigs. Because a primary goal was to test the device, many minor adjustments were made throughout the experiments to optimize the pig’s responses. Thus, these may serve better as proof of concept that pigs can be trained on a task rather than strong baseline data for either task.

#### Experiment 1 – Delay Discounting Task

The goal of this behavior is to assess choice impulsivity [22]. This experiment was performed on the first iteration of the touchscreen device, which ultimately broke. After learning to respond to boxes on the screen, pigs were presented with a magnitude discrimination of two buttons. One delivered 4 pellets (“Large” button), and the other 1 pellet (“Small” button). 6 forced-choice and 6 free-choice trials were given. The first step was to train a magnitude discrimination such that pigs showed preference for the Large button. After that, delays to reinforcement on the Large button were then introduced progressively across the session every 12 trials (0, 5, 10, 20 s). Because pigs rapidly became delay averse, several behavioral manipulations were made to improve stability of choice (described in results). These extended modifications served to provide a long period of assessment for the device.

#### Experiment 2 – Visual Discrimination

The goal of this behavior is to assess the ability to discriminate based on color. This experiment was performed on the second iteration of the touchscreen which is described fully in the methods. After learning to respond to buttons on the screen, pigs were presented with a yellow box on the center of the screen, and after a response, a yellow and blue box on the left and right side of the screen (pseudorandomly presented). Like in experiment 1, multiple behavioral manipulations were performed to improve performance and are described in the results.

## 3. RESULTS

### 3.1 Device Durability

The first iteration of the touchscreen device was ultimately not strong enough and was not described above. An aluminum base and frame mounted a conventional capacitive (home/office grade) touchscreen monitor and was enclosed in a plexiglass covering. Because pigs could get under the plexiglass box, they tended to root at it, and the frame was not heavy enough to prevent this. Ultimately, they broke the screen despite repeated attempts to shape behavior away from such rough interactions. Much of this damage came from lifting the frame and letting it fall, thus it may still be possible to use a capacitive touchscreen in the second iteration, however researchers should anticipate frequent replacements as a common screen protector will not be sufficient long term.

The second iteration, which is fully described in the methods, was much more robust. At the conclusion of the experiment, there was no obvious damage to the device, although pigs did begin to root at it more and additional weight in the bottom may have been beneficial to minimize this. A slightly smaller version of the frame (12U-rack) may have been suitable as well to reduce movement when pushed on and would still accommodate large Yorkshire pigs.

### 3.2 Behavioral Topology of Pigs on the Touchscreen

#### Capacitive vs. Resistive Touchscreen

Version 1 of this device used a capacitive touchscreen (similar to modern smartphones) while version 2 (described in methods) used a resistive touchscreen (similar to ordering kiosks). Pigs learned initial touch responses more rapidly to the capacitive touchscreen, but it is unlikely to hold up to long-term testing. An alternative might be for researchers to swap a capacitive screen in for the initial stages of training, then replace with a resistive screen.

#### Presses on Touchscreen

Overall, pigs were reasonably accurate at pressing buttons. However, it should be noted that the tendency is to press and then swipe upward at a slight diagonal. This resulted in many initial problems as the software was designed to record/act on button *release* rather than press. Even after fixing the buttons, pigs would often have inaccurate responses slightly above the button. For one (male) pig, this resulted in more pronounced swiping behavior. A future option may be to present a slightly larger but invisible response box which extends above the visible button. Care should be taken in considering the layout of buttons in a task.

#### Rooting

Pigs engaged in rooting behavior as noted above. This was largely mitigated in the second version by a frame which they could not get their nose under. However, there were still rooting behaviors present. This seems to be most mitigated by lack of device movement. If an object moves, the pig is more likely to root at it. Smaller/heavier objects are likely to be best for this. For the first experiment, which was performed with a weaker device, rooting behavior was mitigated by instigating a differential reinforcement of other (DRO) behavior schedule during the intertrial intervals. This involved periodically giving marshmallows in other locations of the room to reinforce exploratory behaviors instead of rooting at the screen. In the second version of the device, the pellet dispenser was attached to a remote to reduce rooting toward the screen.

#### Responsiveness to Tones

Tones were added to each response to help pigs discriminate when a response was made. Because there is no tactile feedback on the touchscreen, this helped to distinguish that a press had been recorded. Pigs robustly responded to tone presentation when paired with a reinforcer. A different pitch tone was also used to indicate the start of a trial and was reasonably successful.

#### Frustration and Sensitivity to Reinforcer Magnitude

Pigs were very sensitive to when reinforcement was withheld. This was evident on the few occasions the pellet dispenser failed to deliver: an experimenter would have to give a marshmallow in most cases before pigs would leave the pellet area and re-engage with the touchscreen. Experiment 1 also demonstrated immediate aversion to delayed reinforcement, and similar frustrations were seen on the transition from FR-1 to FR-3 in Experiment 2. Conversely, larger numbers of pellets motivated more rapid re-engagement with the task. Gradually changing response requirements or reinforcer density is recommended.

#### Competing Exploration

All pigs explored extensively during intertrial intervals and often during trials themselves. Items on the walls (e.g., sink, hose) competed for the attention of the pigs. This was more drastic in Experiment 2 which took place in a larger room. The addition of a tone to indicate a trial start helped pigs to orient to the device. Trial durations were increased to allow more time to respond. Intertrial intervals were kept short (<20 s). A reasonably small, plain room with minimal distractions is recommend.

### 3.3 Device Components & Evaluation

#### First Touchscreen Device

The first iteration of our touchscreen broke as described in Device Durability, above. From this, we identified that the standard capacitive (office/home grade) touchscreen was not strong enough. This was largely for two reasons, 1) food was delivered from just under the screen (hopper behind screen and dropped out just in front of screen) and 2) pigs could get under it to lift with their snout. These problems were both changed for the second iteration which is what is fully described in the methods. Other core components, including pellet dispenser, step up/down PCB, and raspberry pi for operation were satisfactory. Supplemental video S1 shows pigs learning to push the buttons on this version.

#### Second Touchscreen Device

Performance of all components of the second iteration were satisfactory and are fully described in the methods. Some initial problems were found with the infrared remote control of the dispenser. Stray infrared signals will be picked up, so care should be taken to monitor which commands are received from other electronics in the area and select input numbers which will not occur from other electronics. It may be advisable to adjust the program such that it periodically turns off all outputs in the absence of a received “on” signal or to use encoding to secure against stray interference. We located the pellet dispenser (and tone generator, and Pi/PCB) on a box placed on top of a sink because it was a convenient method to elevate it above the pigs’ reach. However, a wall-mounted shelf, or 3D-printed hanger would also be sufficient.

#### Extension to Rodents or Other Animals

We plugged the printed PCB and Pi into a Med-Associates rat operant testing chamber and were able to control 28V inputs (two levers, one nose poke) and outputs (pellet dispenser, houselight, levers’ extension, food hopper light) using the standard 7” Raspberry Pi touchscreen. Thus, the same equipment should be capable of running experiments for other animals. However, care should be taken when going across species. For instance, rats and mice do not respond well to a capacitive touchscreen in our experience. This may be why many current commercial solutions use infrared touchscreens in currently available equipment.

### 3.4 Pre-Training and Response Shaping

Pigs were habituated to the researchers through feeding of treats for two days. Pigs were then acclimated to leash walking to the testing room over the course of 2-5 days as described in the methods. The younger, male pigs (experiment 1) took longer, while the female pigs (experiment 2) were able to be put on the leash on the first day.

Pigs were then trained to respond to the touchscreen and gradually shaped to respond to a small, yellow box which moved around the screen. For the male pigs in Experiment 1, seven total sessions were required for them to accurately track the small box on the screen. For the female pigs in Experiment 2, the initial touch response took longer, presumably due to the resistive screen requiring a firm press compared to the capacitive screen in the first device. This was solved by modifying the program to allow any touch to the screen to be immediately reinforced and then by placing ketchup on the screen for one pig. After this, 4-6 sessions were required to shape the response to small boxes. Three sessions additional training was then performed to determine how small of a box the pigs could accurately respond to. Pigs were able to respond to a square occupying as little as 22% of the screen height. Supplemental video S2 shows pigs progressing through the stages of response shaping.

Based on these results, we would recommend researchers habituating pigs in vivarium to experimenter handling, leash, and walking over the course of 5 sessions. The response shaping procedure could likewise be accomplished in 5-7 sessions. Though we tested daily, this timeline could be sped up with multiple sessions per day.

### 3.5 Experiment 1 – Delay Discounting Task

After response shaping, pigs began the magnitude discrimination. Pigs rapidly acquired preference for the Large button across 3-5 sessions. Delays of 0, 5, 10, and 20 s (incremented every 12 trials) were then introduced to the Large button. This caused on increase in initiation latency (Fig 2A), but no large change in omissions (Fig 2B), which held mostly constant across subsequent testing. However, pigs became extremely averse to the delay as indicated by a drop in the Area Under the Curve (AUC) across five sessions (Fig 2C). The AUC represents the proportion of total choices of the large lever (minimum: 0; maximum: 1). By the final session, almost no choice of the Large lever was made, even at 0 s delay. Because pigs were so averse to these longer delays, the Buttons were reversed and delays decreased to 0, 2, 6, and 10 s. However, pigs again rapidly lost preference (5 sessions) for the Large lever, even at the lowest delays. To attempt to fix this, we reversed the Buttons again and implemented another magnitude discrimination. Within 3 sessions pigs strongly preferred the large lever with no delay. We then implemented another set of delays at 0, 1, 2, and 4 s. One pig tolerated this well and the other displayed some discounting but still preference for the large lever. At this point in the experiment, the first iteration of our device broke, and we were unable to further evaluate whether these delays could be titrated out further in a more gradual fashion.

**Figure 2.**
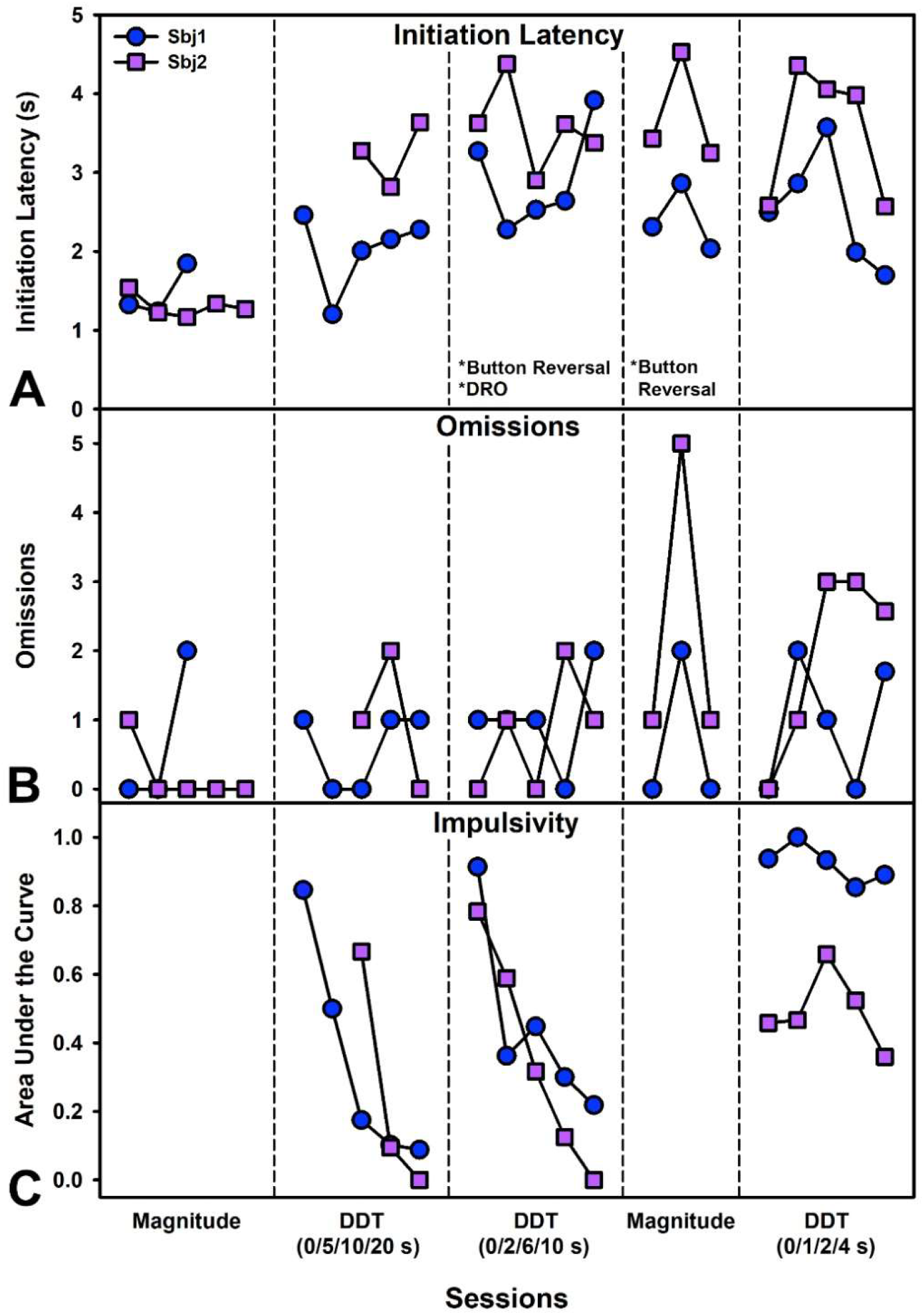
Performance on the Delay Discounting Task (DDT) for two male pigs (Experiment 1). Partitions from left to right indicate performance during a magnitude discrimination (only large versus small buttons, no delays), performance on the DDT with delays of 0, 5, 10, or 20 s (incrementing every 12 trials within the session), performance on the DDT with delays of 0, 2, 6, or 10 s after reversing the levers and giving marshmallows on a differential reinforcement of other (DRO) behavior schedule, performance on a second magnitude discrimination after reversing levers, and finally performance on the DDT with delays of 0, 1, 2 and 4 s. A) Mean latency per session to initiate trials after a tone played indicating button availability. Latencies increased once delays were introduced. B) Total omissions per session. Omissions remained relatively low throughout testing. C) Area under the discounting curve which indicates the proportion of large/delayed/self-controlled choices made. Pigs made progressively more impulsive choices during the first set of delays but performed better at the low delays trained at the end.

### 3.6 Experiment 2 – Color Discrimination

After the response shaping, pigs began the color discrimination. Choices of the yellow button were reinforced (blue was the comparison color). During the first phase free choices were made throughout the session. During the second phase, a correction trial was implemented such that if an incorrect choice was made, the same trial was immediately represented but the incorrect option could not be chosen (box appeared but as inactive). Once pigs were accurate at this, the third phase presented a more difficult conditional discrimination where a single-colored button (green or blue) was presented in the center and then only choices of that color were reinforced. As performance degraded in the third phase, an FR-3 requirement was put on the center (comparison) button and also on the choice buttons to try and increase salience. Pigs initiated trials within approximately 8 s, but latencies to start increased as performance went down in the conditional discrimination (Fig 3A). Similarly, number of trials completed was similar for each pig across all conditions until performance went down (Fig 3B). Omissions were initially high for one pig but remained low throughout the rest of testing (Fig 3C). Accuracy on the task was very low until the correction trial was implemented and then rapidly increased over 1-2 sessions. Performance dropped sharply during the conditional discrimination, suggesting additional training would be needed. The FR-3 requirement was not sufficient to rescue accuracy (Fig 3D). Pigs also had a tendency toward side biases (Fig 3E) and a bias toward the green color once the conditional discrimination was implemented (Fig 3F). Supplemental video S3 shows pigs performing the discrimination task.

**Figure 3.**
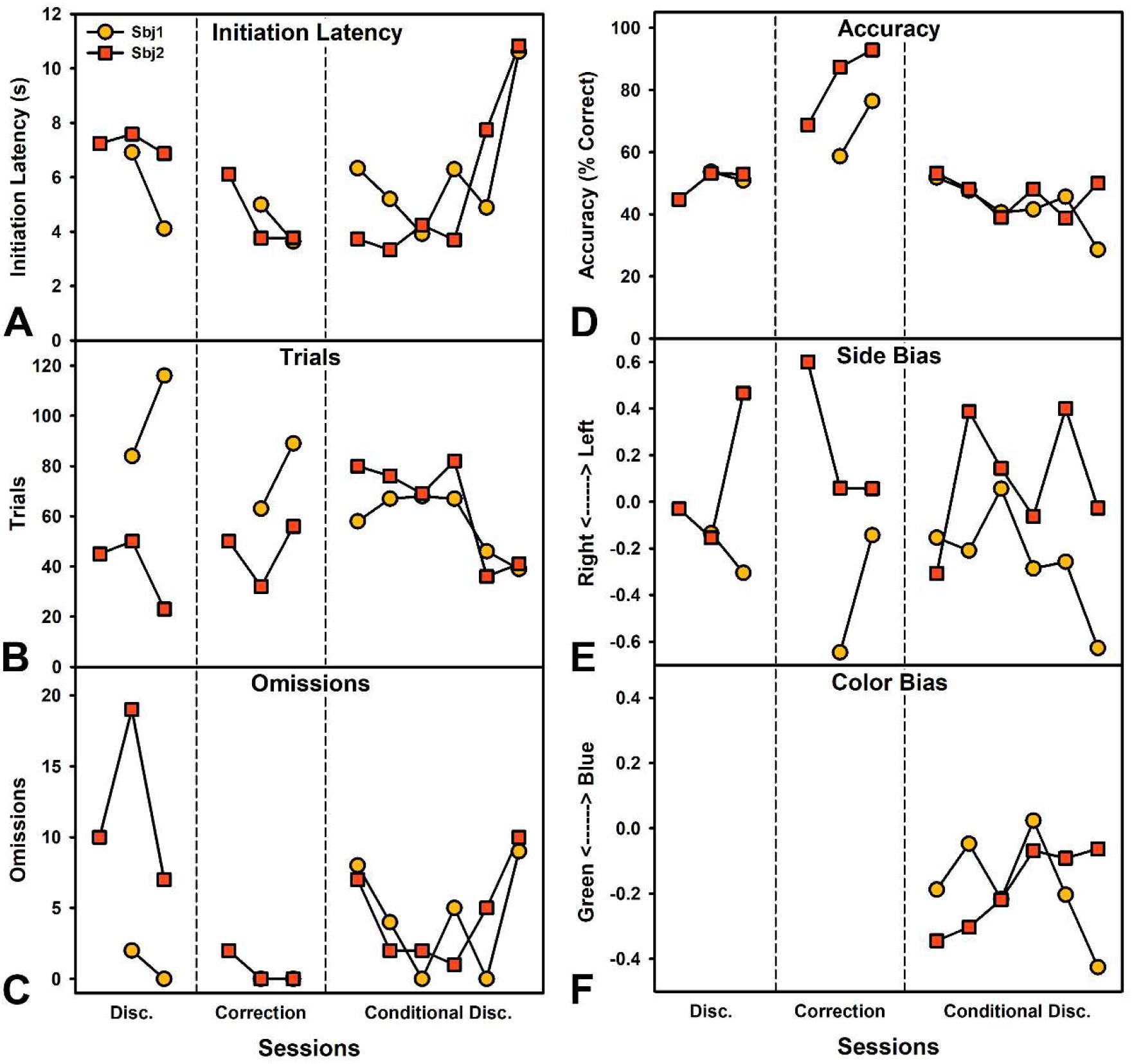
Performance on a Color Discrimination task for two female pigs (Experiment 2). Partitions from left to right indicate performance during an initial discrimination (yellow choices reinforced), the same discrimination but with correction trials which forces choice of the correct option after an incorrect choice, and finally a conditional discrimination where a blue or green box was shown in the middle and then two choice boxes presented and only choices of the presented color were reinforced. A) Mean latency per session to initiate trials after a tone played indicating button availability. Latencies increased as performance worsened under the conditional discrimination. B) Total trials completed per session. Trials remained relatively high throughout. C) Total omissions per session. Omissions for one pig were large during initial performance, and then somewhat variable under the conditional discrimination. D) Percent correct choices. Pigs performed poorly until the correction trials were implemented and then rapidly improved. Conditional discrimination performance was very poor. E) Preference for left versus right (−1 to 1 scale with 0 indicating no preference). Pigs exhibited moderate side biases throughout testing. F) Preference for green versus blue color (−1 to 1 scale with 0 indicating no preference). Pigs showed a strong preference toward green, which may be more similar to the previously-reinforced yellow.

## 4. DISCUSSION

In the current protocol, we provide a detailed description for constructing an operant touchscreen for pig behavioral testing and assessed two behavioral tasks, highlighting the utility of such devices. Previously, pig behavior had largely been limited to relatively simplistic measures or required room-sized equipment. The current touchscreen device is only limited by the time constraints of the researcher to train. Although pigs are not a common laboratory species for all disciplines, there are several key research areas that could benefit from such a device.

The fields with the strongest integration of pigs are those of pharmacology, cardiology, cancer, and vision. Pigs have long been used for safety testing of drugs and represented a critical translational step from rodent safety and efficacy studies [23]. For drugs affecting peripheral physiology, the pig has made a strong model due to the many physiological similarities. However, for psychoactive drugs, researchers have historically relied on monkeys to obtain both efficacy data and safety data. With a touchscreen device, pigs could become a viable alternative for testing psychotherapeutics. In vision research, pigs were classically used from the 1960s through the 1980s [24], and in recent years there has been a resurgence of interest in pigs to study retinal degeneration [25]. A recent study established an obstacle course behavioral test to assay gross visual impairment [26] but a touchscreen assay could provide much more detail on the nature and progression of deficits by systematically manipulating the salience of stimuli.

In the field of TBI, pigs have become much more common in recent years [27]. Their large and gyrencephalic brains make them ideal specimens for examining pathophysiology associated with rotational damage and unique cortical damage which cannot be observed in rodents. However, functional assessments have been much more limited. Now, functional assessments with great relevance to brain injury can be used. Behavioral flexibility, attentional impairments, and impulsivity are all symptoms of brain injury in patients and could not be readily assessed without a device such as this. In the current paper, we report an example of delay discounting, which could be used to measure impulsivity in a pig model of brain injury. This would provide crucial data about a relatively common psychiatric outcome of TBI [28]. More robust and extended behavioral batteries will need to be developed to meet the needs of the TBI field. In particular, both assays that are rapid (for acute studies) and those that can hold up to extended, repeat testing (for chronic studies) will be important.

Still other fields are being shaped by the recent advent of transgenic minipigs. The current device could be immediately adapted to the minipig by dropping the screen down lower on the frame and would require no other changes. It could feasibly be used in labs which work in both mini- and full-size species. Perhaps most immediately relevant to studying behavior are the transgenic Göttingen minipigs for HD [5] and for AD [6]. Memory tasks or others could be easily programmed for assessment. However, these same pigs likely have even broader applications. Systems for rapid gene editing (e.g., CRISPR) are now being used to reduce the cost of generating a unique transgenic minipig for a given question of interest [29, 30]. Thus, a device such as this touchscreen, with the flexibility to design multiple behavioral assessments provides great utility for these research questions. A small but encouraging literature exists describing these types of assessments in pigs. Researchers have evaluated impulsivity, memory, and decision-making [12–17] using non-touch operant devices.

While there are numerous fields that could benefit from adopting behavioral testing using an apparatus such as the one described here, there are also several limitations to consider. First, while we have described a method for constructing a device with relative ease, individual researchers will still need to program relevant behavioral tests for their questions of interest. Perhaps more challenging is the need for behavioral expertise. For a physiology researcher with no background, it may be difficult to adapt to the needs of behavioral study (e.g., training time). However, colleagues in psychology and neuroscience departments may be readily available to provide such expertise. Despite stark differences in rat or human behaviors, many such researchers regularly program tasks such as the ones described in this paper. Our own lab is otherwise focused on rat behavior, but were able to design and program these tests for pigs.

Researchers must also be ready to recognize and adapt to the limitations of pig behavior. As an example, in the current study, we quickly determined that the method by which a pig pushed a button on a screen was not congruent with the default expectations for the software. Pigs tend to swipe in an upward direction as opposed to the built-in function expecting to execute a command on release (e.g., mouse click or finger tap). Thus, pigs would start on a button and swipe off it before release. This was easily mitigated by processing a response on depression of a button rather than release. Further testing may identify whether a larger or taller response box to record the press might be helpful for this same problem. Similarly, we had challenges with rooting behaviors, and indeed our first device was broken by the pigs. Design refinements to reduce movement and access to areas under the screen as well as move the dispenser solved some of these problems. However, rooting is a problem that was noted as early as 1961 in a classic article titled “The Misbehavior of Organisms”. This work noted (with pigs as one example) that there was often drift toward instinctual responses (e.g., rooting) after extended training with food reinforcement.

The current paper represents a start to automate and extend operant testing in pigs. There is still more work to be done to optimize behavioral training regimens which will reflect the needs of various fields. For example, rapid acquisition tasks for short time frames (e.g., <10 days) versus more extended in-depth repeated measurements of stable trait behaviors. Researchers may want to extend on this and integrate other peripheral elements or input devices. For instance, a physical button, a foot lever, or a lever which could be manipulated with the mouth can all be integrated using the device described here. It could even be taken into rats or other species if a suitable response device can be found (likely an infrared touchscreen instead of resistive or capacitive). Multiple reinforcer types could be used with a behavioral economics approach to tease out subtle aspects of preference and motivation. Or two screens could be used alongside one another (controlled by one Pi or coordinating multiple) to provide stronger spatial separation of choice boxes. The shaping program described in this paper will be available on the corresponding author’s GitHub. Additional updates to this project and programs will be made available as they are developed.

## Acknowledgements

Special thanks to Gregory Lusk and Doug Mathess who helped with initial design aspects. Thanks to Injury and Recovery Lab members who assisted with behavioral testing. Funding for this project was provided by West Virginia University and Ohio State University.

## Supplemental Material

Three videos are available which show the first device (S1), acquisition of pressing behavior (S2), and the color discrimination (S3). A schematic for the PCB in provided as S4. A program and other online resources are provided on the corresponding author’s GitHub repository, https://github.com/VonderHaarLab/.

## Data Availability Statement

Resources to create reproduce this device are available on the corresponding author’s GitHub repository. Raw data will be made available upon request.

